# Biogeography of the savanna-like vegetation in hot dry valleys in southwestern China with reference to their floristic origin and evolution

**DOI:** 10.1101/402545

**Authors:** Hua Zhu, Lichun Yan

## Abstract

Savanna-like vegetation and dry thickets occur in hot dry valleys across southwestern China. Here, the flora and biogeography of these vegetations are studied. Native seed plants of 3,217 species from 1,038 genera in 163 families are recorded from the hot dry valleys in SW China. The biogeographical elements with a tropical distribution contribute 57.18%, but the ones with a temperate distribution contribute 36.45% of the total genera of the flora. This shows that the flora has proliferated by temperate elements via their evolution, although the flora occur in tropical habitats in the hot dry valleys. Floristic divergence across these hot dry valleys is obvious. The floras in the Yuanjiang (the upper reaches of the Red River) and the Nujiang (the upper reaches of the Salween River) valleys are dominated by tropical elements (77.26% and 74.49 of the total genera, respectively), but the flora of the Jinshajiang (the upper reaches of the Yangtze River) valley is composed of half tropical (47.27%) and half temperate (44.96%) genera. Regarding floristic similarities, the Jinshajiang shows the highest similarity to the Yuanjiang although these river valleys are located a great distance from each other. Our results could be well explained from the geological events since the Cenozoic, such as the uplift of Himalayas, the extrusion of Indochina, the river capture of the Jinshajiang separating from the Yuanjiang, and the northward movement of the Burma Plate. Further floristic comparison between the flora in hot dry valleys of SW China and southern Africa supports the consideration that the flora of savanna-like vegetations of SW China could have floristic affinity to African savannas over the course of its evolutionary history by the Indian Plate from southern Africa colliding with Eurasia in the Cenozoic.

## Introduction

The savanna-like vegetation or dry thickets, including woodland and dry thorny shrubs, occur in hot dry valleys and some dryer habitats in southeastern Asia [1]. Scholes and Archer clarified that savannas are defined on the basis of both ecological characteristics and climatic attributes, and the common characteristic of all savannas is the coexistence of woody and herbaceous vegetation in regions where seasonality is controlled by distinct dry seasons rather than by cold [2]. The savanna-like vegetation in hot dry valleys in southwestern China meets this definition.

The savanna-like vegetations in southwestern (SW) China are distributed mainly in Yunnan Province in the deep, hot, and dry valleys of several rivers. Yunnan Province is between 21°09′ to 29°15′N and 97°32′ to 106°12′E and borders Myanmar to the west and Laos PDR and Vietnam to the south and southeast, respectively (Fig. 1). It has a mountainous topography with the mountain ridges generally running in a north-south direction, decreasing in elevation southward. Its elevation ranges from 76.4 m at the lowest valley bottom in the southeast (Red River) to 6,740 m at the highest mountain summit in the northwest. Yunnan is extremely diverse in habitats and topography. It commonly has tropical dry climates in deep valleys below 1,000 m with an annual mean temperature of approximately 21–24 °C and an annual precipitation of 600–800 mm, due to the foehn effect [3]. Yunnan is a region with tropical areas at the horizontal base because almost all areas of lower elevation are tropical in nature regardless of their latitudinal location. Large and continuous savanna-like vegetations in Yunnan occur mainly along the Jinshajiang (the upper reaches of the Yangtze River), the Yuanjiang (the upper reaches of the Red River), and the lower reaches of the Nujiang (the upper reaches of the Salween River) (Fig. 2).

**Figure 1.**
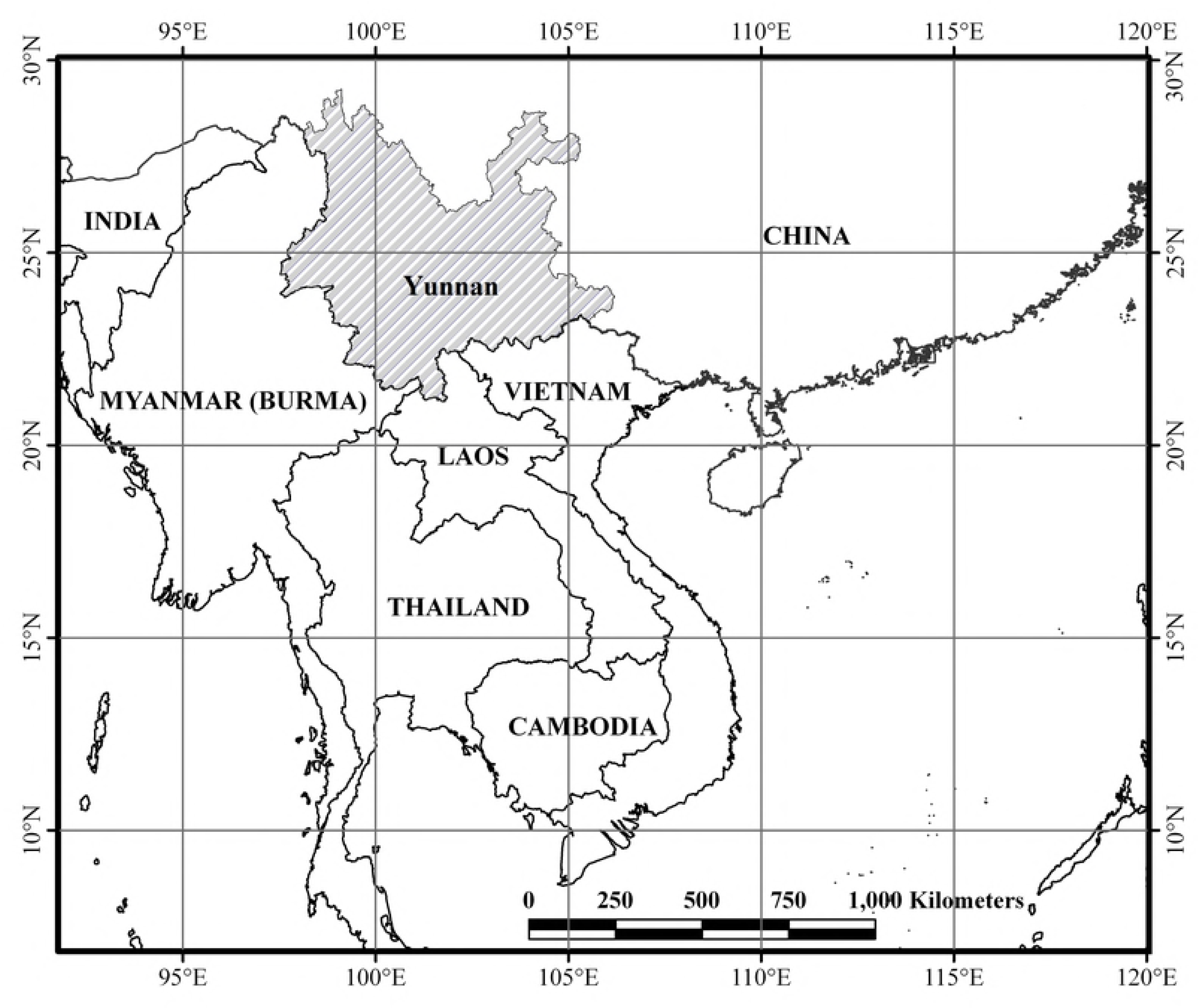
Geographical location of Yunnan Province, southwestern China.

**Figure 2.**
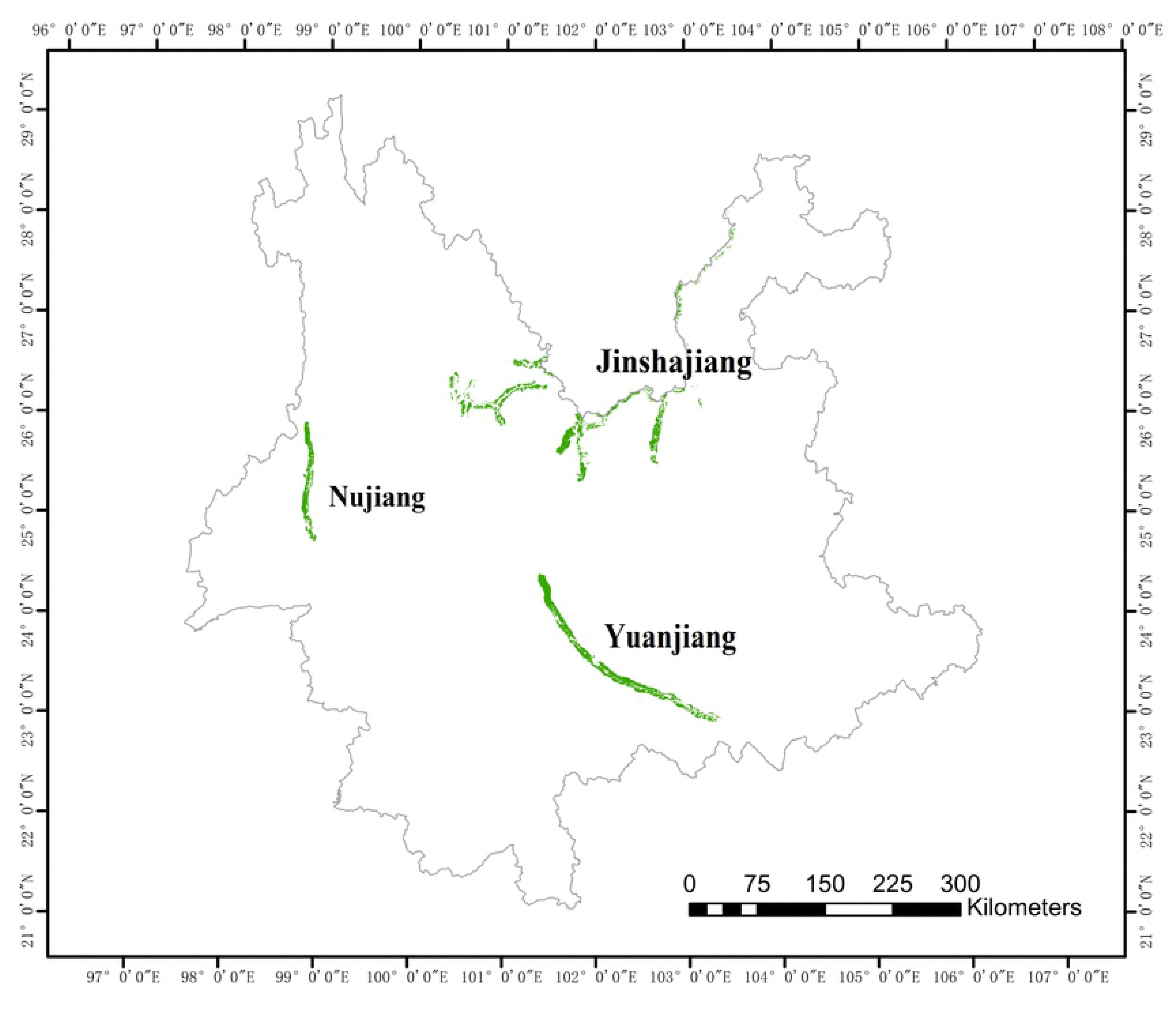
Distribution of the savanna-like vegetations in the hot dry valleys of Yunnan Province in southwestern China.

The savanna-like vegetations in SW China are dominated mainly by herbaceous plants (grasses), dotted with sparse trees, and have some sites with shrubs [4]. Dry thorny shrubs are also present [5]. They are dominated by grass species, such as *Heteropogon contortus* and *Cymbopogon citratus*, shrubs, such as *Dodonaea viscose*, *Acacia farnesiana*, *Woodfordia fruticosa* and *Euphorbia royleana*, and trees, such as *Bombax ceiba*, *Lannea coromandelica, Phyllanthus emblica*, and *Buchanania latifolia*. The savanna-like vegetations look similar to the savannas in Indian-Myanmar and Africa in physiognomy (see photo1 and 2) and show floristic affinity to African savannas with some species, such as *Woodfordia fruticosa* and *Calotropis gigantea*. For example, the genus *Woodfordia* includes two species: one in Africa and the Arabian Peninsula, and the other in SE Asia and SW China [6]; the genus *Calotropis* includes three species in northern Africa, Arabia, and tropical Asia. The two genera are characteristic elements of African savannas. *Woodfordia fruticosa* and *Calotropis gigantea* are native to the savannas in Yunnan Province, and these species offer clues for the floristic affinities of the savannas.

The floras in hot dry valleys of SW China were primarily studied at the local scale [5-9]. Later the floristic composition and characteristics of the vegetation in tropical and subtropical dry valleys, including savanna-like vegetation, were also studied [10]. However, a complete floristic survey on the savanna-like vegetation in hot dry valleys of SW China are lacking. Numerous studies have been conducted on savannas worldwide. Most of these studies have focused on ecology such as studies conducted in African savannas [11] and in tropical America [12]. Floristic and biogeographical studies on savannas are limited. A biogeographical study based on the complete floristic inventory of savanna-like vegetations in SW China is important to understand its biodiversity, biogeography, and floristic affinity.

The origin and evolution of the flora in the hot dry valleys in SW China have been of interest to scientists for decades. They are related to the geological history of the region, such as the collision between India and Eurasia, the uplift of the Himalayas [13], and river capture events [14, 15]. It has been suggested that the modern rivers draining in the plateau margin of SW China, i.e., the Jinshajiang, Nujiang, and Lancanjiang (the upper reaches of the Mekong River), were once tributaries to a single, southward flowing system—the paleo-Red River, which drained into the South China Sea. Disruption of the paleo-drainage occurred by river capture and reversal prior to or coeval with the initiation of the Miocene uplift of the Himalayas [15]. Biogeographical studies on the flora in hot dry valleys of these rivers could provide help to the geological events in SW China.

The aim of this article is to understand the biodiversity, biogeography, and floristic affinity of the savanna-like vegetations in SW China, and to provide evidence for the geological events in the region.

## Materials and methods

We compiled floristic inventories of the savanna-like vegetations in the hot dry valleys of the reaches of the three main rivers in Yunnan Province, SW China: the Jinshajiang, Yuanjiang, and Nujiang, based on intensive fieldwork and specimen identifications, as well as data collection from the herbariums of Kunming Institute of Botany and Xishuangbanna Tropical Botanical Garden. We obtained a complete native species list of seed plants that is composed of 3,217 species from 1,038 genera in 163 families. Family circumscription follows APG III [16, 17] and species names were rectified following the nomenclature and classification presented in the w^3^TROPICOS database housed at the Missouri Botanical Garden (http://mobot.mobot.org/W3T/Search/vast.html). The inventory information is presented in supplementary material S1(Appendix 1). Patterns of seed plant distribution were quantified at the generic level mainly based on Wu [18] and Wu et al. [19], and at the family level following Wu et al. [20]. We classified geographical elements on distribution types at family and generic levels as: cosmopolitan, pantropic, tropical Asia and tropical America disjunct, Old World tropics, tropical Asia to tropical Australia, tropical Asia to tropical Africa, tropical Asia, north temperate, East Asia and North America disjunct, Old World temperate, temperate Asia, Mediterranean region and west to central Asia, central Asia, East Asia, and endemic to China. We analyzed the floristic and geographical attributes of the native seed plants. The flora in hot dry valleys was compared with the floras of the neighboring regions including Myanmar to the west and Guangxi Province, China to the east as well as southern Africa. The geological history and river capture events that occurred in SW China are also discussed from our biogeographical studies on the savannas.

## Results

### Floristic composition

The native seed plants of 3,217 species from 1,038 genera in 163 families were recorded from the hot dry valleys of the three main rivers (Jinshajiang, Yuanjiang, and Nujiang) in SW China. Families with more than 100 species include Fabaceae (68 genera/307 species), Poaceae (96/253), Asteraceae (77/212), Lamiaceae (38/175), and Rosaceae (32/167). Other species-rich families are Rubiaceae (27/83), Euphorbiaceae (26/80), and Orchidaceae (40/68) (Table 1). The most species-rich families (with more than 100 species each) also have an almost cosmopolitan distribution. However, in the top 20 species-rich families of the savanna flora, six families have pantropic distribution, and 13 families have cosmopolitan distribution (Table 1).

**Table 1.**
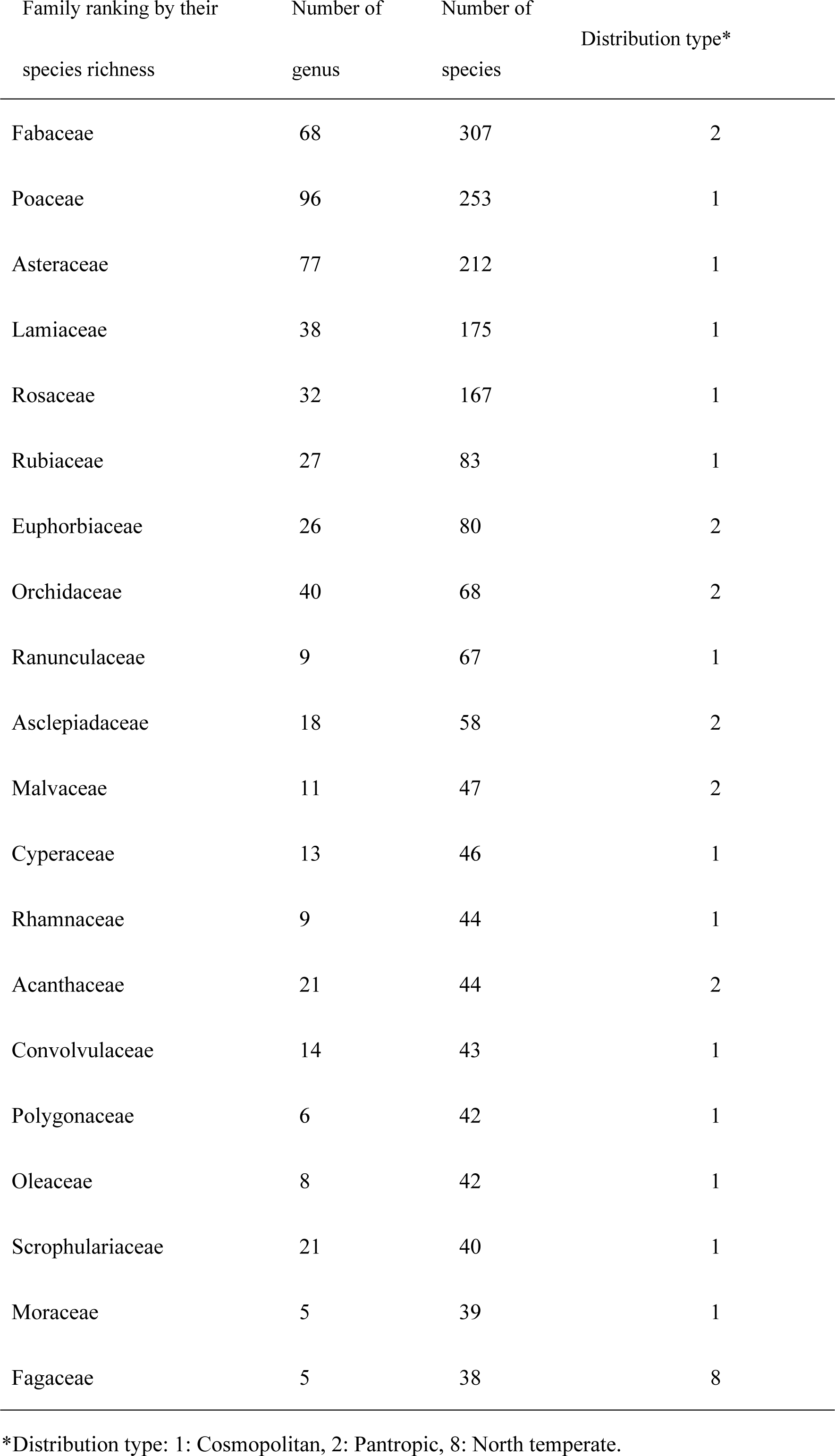
Dominant (top 20) families in species richness with their distribution in the flora in hot dry valleys in SW China

| Family ranking by their species richness | Number of genus | Number of species | Distribution type* |
| --- | --- | --- | --- |
| Fabaceae | 68 | 307 | 2 |
| Poaceae | 96 | 253 | 1 |
| Asteraceae | 77 | 212 | 1 |
| Lamiaceae | 38 | 175 | 1 |
| Rosaceae | 32 | 167 | 1 |
| Rubiaceae | 27 | 83 | 1 |
| Euphorbiaceae | 26 | 80 | 2 |
| Orchidaceae | 40 | 68 | 2 |
| Ranunculaceae | 9 | 67 | 1 |
| Asclepiadaceae | 18 | 58 | 2 |
| Malvaceae | 11 | 47 | 2 |
| Cyperaceae | 13 | 46 | 1 |
| Rhamnaceae | 9 | 44 | 1 |
| Acanthaceae | 21 | 44 | 2 |
| Convolvulaceae | 14 | 43 | 1 |
| Polygonaceae | 6 | 42 | 1 |
| Oleaceae | 8 | 42 | 1 |
| Scrophulariaceae | 21 | 40 | 1 |
| Moraceae | 5 | 39 | 1 |
| Fagaceae | 5 | 38 | 8 |

\*Distribution type: 1: Cosmopolitan, 2: Pantropic, 8: North temperate.

At the generic level, *Clematis* (37 species), is the most species-rich genus, followed by *Indigofera* (34 species), *Ficus* (29), *Rosa* (29), *Artemisia* (28), *Isodon* (27), *Cotoneaster* (24), *Dioscorea* (22), *Rubus* (22), and *Elsholtzia* (21) (Table 2). In these species-rich genera, their distributions vary compared to the species-rich families. The top 20 species-rich genera include seven distributions: five genera with pantropic distribution, five with north temperate distribution, four with cosmopolitan distribution, and three with Old World temperate distribution.

**Table 2.** Dominant (top 20) genera in species richness with their distribution in the flora in hot dry valleys in SW China

| Genera ranking by their<br>species richness | Number of<br>species | Distribution type* |
| --- | --- | --- |
| *Clematis* | 37 | 1 |
| *Indigofera* | 34 | 2 |
| *Ficus* | 29 | 2 |
| *Rosa* | 29 | 8 |
| *Artemisia* | 28 | 8 |
| *Isodon* | 27 | 6 |
| *Cotoneaster* | 24 | 10 |
| *Dioscorea* | 22 | 2 |
| *Rubus* | 22 | 1 |
| *Elsholtzia* | 21 | 10 |
| *Polygonum* | 21 | 1 |
| *Spiraea* | 21 | 8 |
| *Cynanchum* | 20 | 10 |
| *Desmodium* | 18 | 9 |
| *Quercus* | 18 | 8 |
| *Astragalus* | 17 | 1 |
| *Jasminum* | 17 | 2 |
| *Silene* | 17 | 8 |
| *Smilax* | 17 | 2 |
| *Campylotropis* | 16 | 7 |

\*Distribution type: 1: Cosmopolitan, 2: Pantropic, 6: Tropical Asia to Tropical Africa, 7: Tropical Asia (Indo-Malesia), 8: North temperate, 9: E. Asia and N. America disjuncted, 10: Old World Temperate.

### Geographical elements

Out of the 163 families, the ones with a tropical distribution contribute 51.23% including those with a pantropic distribution (40.12%), such as Acanthaceae, Anacardiaceae, Annonaceae, Apocynaceae, Asclepiadaceae, Bignoniaceae, Burseraceae, Combretaceae, Ebenaceae, Euphorbiaceae, and Fabaceae, contributing the greatest ratio. The cosmopolitan families contribute 26.54% of the total families, such as Asteraceae, Lamiaceae, and Poaceae. Families with mainly temperate distributions contribute 22.22% of the total flora including Caprifoliaceae, Fagaceae, and Liliaceae (Table 3).

**Table 3.** Geographical elements of seed plants at family level in the flora in hot dry valleys in SW China

| Geographical elements of family | No. of family | % |
| --- | --- | --- |
| 1 Cosmopolitan | 43 | 26.54 |
| 2 Pantropic | 65 | 40.12 |
| 3 Tropical Asia and Tropical America disjuncted | 9 | 5.56 |
| 4 Old World Tropics | 5 | 3.09 |
| 5 Tropical Asia to Tropical Australasia | 3 | 1.85 |
| 6 Tropical Asia to Tropical Africa | 0 | 0.00 |
| 7 Tropical Asia (Indo-Malesia) | 1 | 0.62 |
| 2–7 (Total tropical elements) | (83) | (51.23) |
| 8 North Temperate | 27 | 16.67 |
| 9 E. Asia and N. America disjuncted | 4 | 2.47 |
| 10 Old World Temperate | 1 | 0.62 |
| 11 Temperate Asia | 0 | 0.00 |
| 12 Mediterranean, West Asia to Central Asia | 1 | 0.62 |
| 13 Central Asia | 0 | 0.00 |
| 14 East Asia | 3 | 1.85 |
| 15 Endemic to China | 0 | 0.00 |
| 8–15 (Total temperate elements) | (36) | (22.22) |
| Total | 162 | 100.00 |

Patterns of seed plant distribution of the flora at the generic level are given in Table 4. The genera with a tropical distribution contribute the greatest ratio, up to 57.18% of the total genera, including genera with pantropic distribution (20.83% of the total sum), such as *Acacia, Bauhinia, Dioscorea, Ficus, Indigofera, Jasminum, Phyllanthus, Smilax*, and *Vitex*. The genera with a tropical Asian distribution, such as *Campylotropis, Pueraria, Engelhardia*, and *Murraya*, contribute 13.98%. The genera with Old World tropical distribution, such as *Cotoneaster, Duhaldea, Elsholtzia*, and *Fagopyrum*, contribute 8.29%. Genera with tropical Asian to tropical African distribution include *Arthraxon, Bombax, Cymbopogon, Flacourtia, Isodon, Miscanthus*, and *Myrsine*, contributing 6.17%.

**Table 4.**
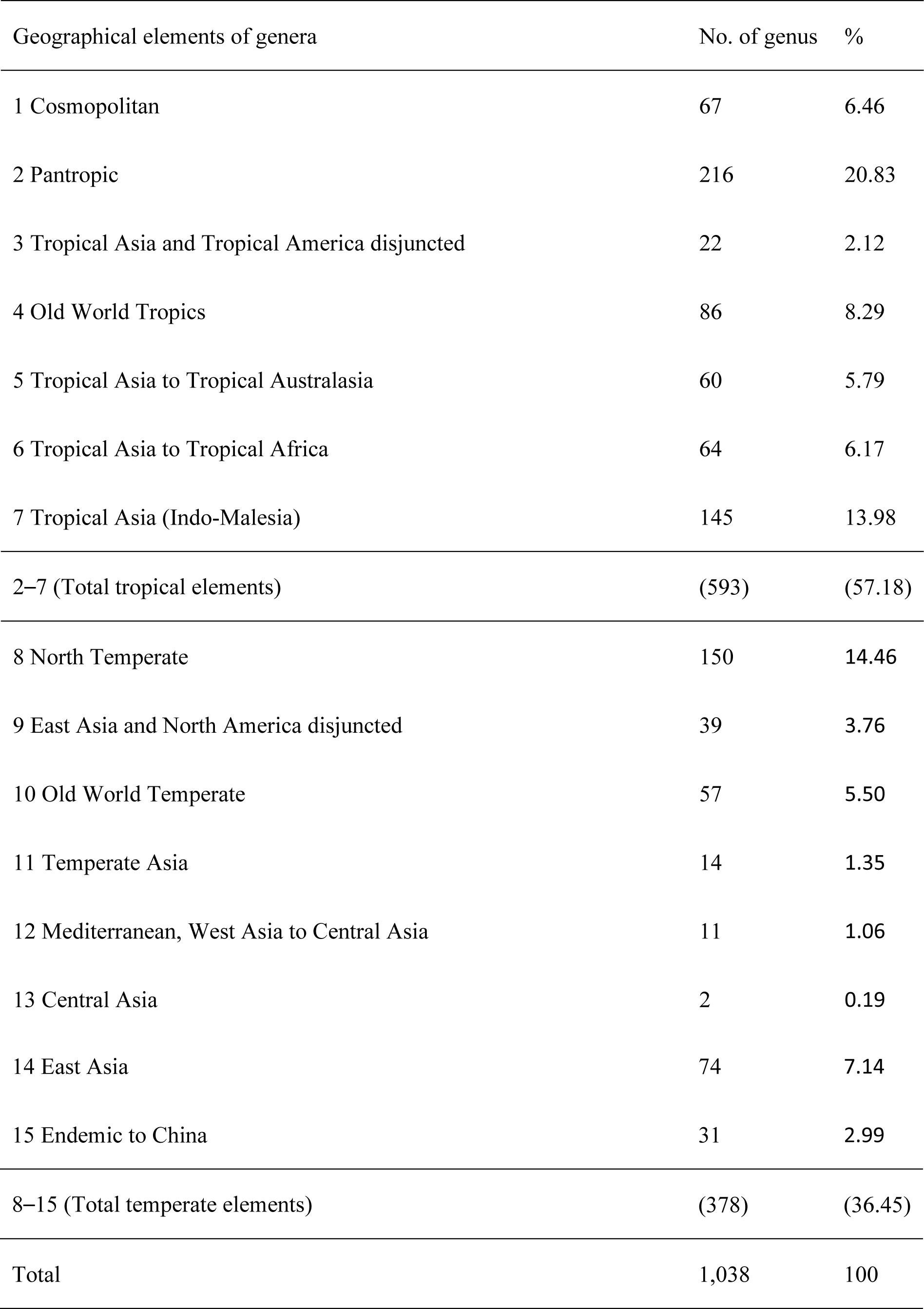
Geographical elements of seed plants at generic level in the flora in hot dry valleys in SW China

| Geographical elements of genera | No. of genus | % |
| --- | --- | --- |
| 1 Cosmopolitan | 67 | 6.46 |
| 2 Pantropic | 216 | 20.83 |
| 3 Tropical Asia and Tropical America disjuncted | 22 | 2.12 |
| 4 Old World Tropics | 86 | 8.29 |
| 5 Tropical Asia to Tropical Australasia | 60 | 5.79 |
| 6 Tropical Asia to Tropical Africa | 64 | 6.17 |
| 7 Tropical Asia (Indo-Malesia) | 145 | 13.98 |
| 2–7 (Total tropical elements) | (593) | (57.18) |
| 8 North Temperate | 150 | 14.46 |
| 9 East Asia and North America disjuncted | 39 | 3.76 |
| 10 Old World Temperate | 57 | 5.50 |
| 11 Temperate Asia | 14 | 1.35 |
| 12 Mediterranean, West Asia to Central Asia | 11 | 1.06 |
| 13 Central Asia | 2 | 0.19 |
| 14 East Asia | 74 | 7.14 |
| 15 Endemic to China | 31 | 2.99 |
| 8–15 (Total temperate elements) | (378) | (36.45) |
| Total | 1,038 | 100 |

The genera with a temperate distribution contribute 36.45% of the flora including the genera with a northern temperate distribution (14.46%), such as *Fraxinus, Lilium, Lonicera*, *Rosa*, and *Spiraea*, and the genera with an East Asian distribution (7.14%), such as *Caryopteris, Chelonopsis, Codonopsis*, and *Leptodermis*. Genera endemic to China contribute 2.99% with 31 genera, such as *Archiserratula, Biondia, Pterygiella, Ostryopsis, Trailliaedoxa*, and *Tsaiodendron.*

### Divergence of the flora across the hot dry valleys of SW China

The three floras in hot dry valleys of the Yuanjiang, Nujiang, and Jinshajiang are compared. The floristic similarities at the family, generic, and specific levels are given in Table 5. At the family level, the three floras are almost the same and the similarity coefficient is more than 87%. At the generic level, the highest similarity is between the Yuanjiang and Jinshajiang (73.84%), and all have a similarity of more than 62%, showing that close floristic affinity remains. However, at the specific level, the similarity coefficients are generally low, with the highest (53.76%) between the Yuanjiang and Jinshajiang, and the lowest (35.03%) between the Yuanjiang and Nujiang. Obviously, the flora of the hot dry valleys in SW China have diverged at the specific level.

**Table 5.**
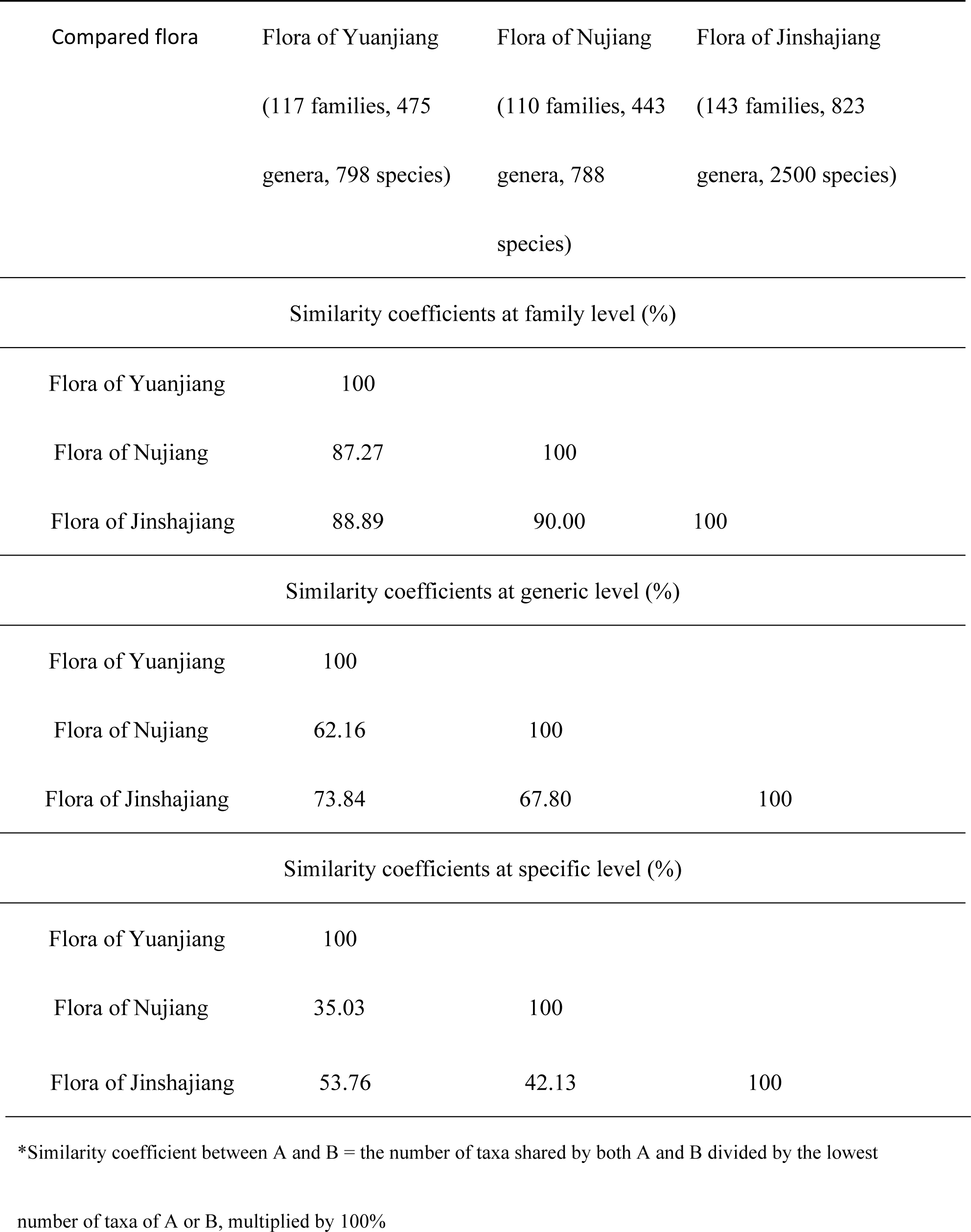
Comparison of floristic similarities at the family, generic and specific levels between the floras of Yuanjiang, Nujiang and Jinshajiang in hot dry valleys in SW China

Geographical elements at the generic level of the three regional floras (Yuanjiang, Nujiang, and Jinshajiang) are given in Table 6. The floras of the Yuanjiang and Nujiang are dominated by tropical elements (77.26% and 74.49 of the total genera, respectively), illustrating their tropical nature. The flora of the Jinshajiang has a similar tropical (47.27%) to temperate (44.96 %) ratio. Further, the floras of the Yuanjiang and Nujiang have more tropical Asia elements (16.42% and 17.38%, respectively) and tropical Asia to tropical Australia (6.53% and 7.90%, respectively) elements. The flora of the Jinshajiang has more north temperate (18.23%) and East Asia (8.14%) elements. These divergences between the three river valleys could be explained by geological history of these regions, although the climates are very similar.

**Table 6.**
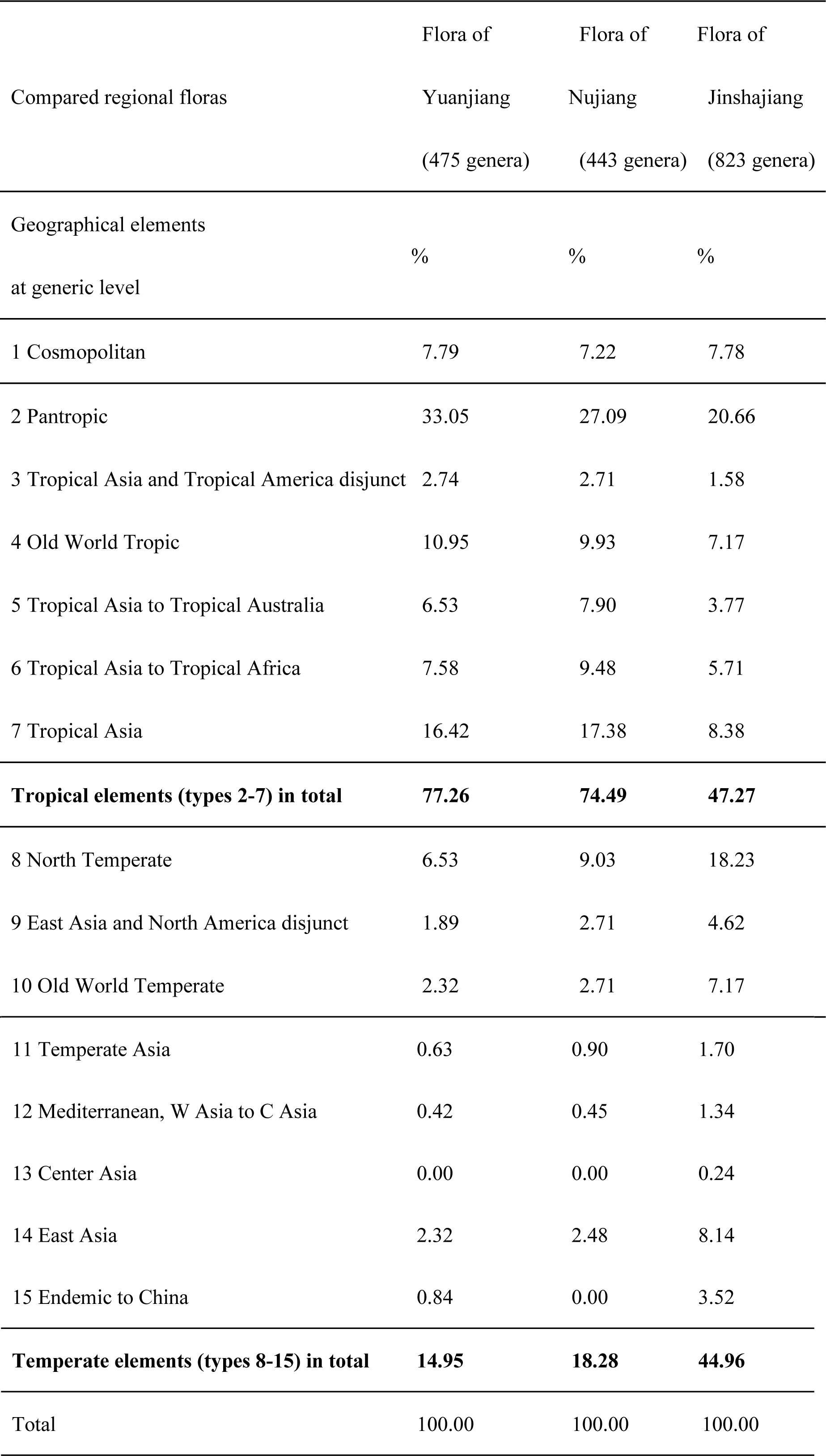
Comparison of geographical elements of seed plants at the generic level between the three floras in hot dry valleys in SW China

## Discussion

In the flora in hot dry valleys of SW China, species-rich families are composed of larger families with cosmopolitan and pantropic distributions. The species-rich genera are those with various distributions, except pantropic and cosmopolitan distribution; genera with temperate distributions appear, especially genera with primarily north temperate and Old World temperate distributions. This is the common scenario for flora from Yunnan Province and SW China [21, 22]. For the total flora, the families with tropical distribution contribute 51.23%, including those with a pantropic distribution (40.12%), making up the highest ratio, and the families with temperate distribution contribute 22.22% of the flora (Table 3). This illustrates that the flora of the studied savanna-like vegetations is fundamentally tropical in nature.

At the generic level, those with a tropical distribution contribute 57.18% of the total genera, including the genera of pantropic distribution, consisting of 20.83%, the highest ratio of all geographical elements. The genera with a tropical Asian distribution contribute 13.98%. The genera with a temperate distribution contribute 36.45% of the flora, including the genera with northern temperate distribution (14.46%) and East Asian distribution (7.14%) (Table 4). Although tropical elements dominate the flora at the generic level, other geographical elements, such as northern temperate and East Asian distributions, are manifested. Geographical elements at the generic level further reveal that although the savanna-like vegetations occur in tropical habitats of the hot dry valleys, they have proliferated by temperate elements via their evolution, which could be due to the geological history of these regions.

Southwest China has faced the upheaval of geoblocks and topographical changes since the Cenozoic. Southwest China and continental Southeast Asia are composed of geoblocks bounded by suture zones [23]. Yunnan Province was formed from the Indochina block or terrane, South China block, Shan-Thai terrane, and West Burma block [24]. The Himalayas formed and were quick uplifted with the collision between India and Eurasia, which began in the early Cenozoic [13]. With the uplift of the Himalayas and together with the extrusion of Indochina [25-29], these deep valleys in SW China formed with a hot dry climate, producing the savanna-like vegetations. In the Jinshajiang, river capture occurred concurrently with the uplift of the Himalayas [15]. These historical events could influence the divergence of the flora in hot dry valleys. Table 5 shows that the floras in the three hot dry valleys included in this study have similar family and generic composition, but they have low similarity coefficients in species composition, indicating that they have diverged mainly at the specific level. It is notable that the floristic similarities at the generic and specific levels are higher between the Yuanjiang and Jinshajiang, although they are distant from each other. Comparison of the geographical elements at the generic level of the three floras in the hot dry valleys show that the floras of the Yuanjiang and Nujiang are dominated by tropical elements (77.26% and 74.49% of the total genera, respectively), but the flora of the Jinshajiang is composed of half tropical (47.27%) and half temperate (44.96 %) elements (Table 6). These differences can be explained by geological history. There is evidence that the Jinshajiang was once a tributary to the paleo-Red River, i.e., the present Yuanjiang. Disruption of the paleo-drainage occurred by river capture and reversal prior to or coeval with the initiation of the Miocene uplift of the Himalayas [15], which led to the separation of the Jinshajiang from the Yuanjiang. *Trailliaedoxa* is an endemic, monotypic genus with a small population that was previously found only in the hot dry valley of the Jinshajiang [30]. During our fieldwork, we found a large population of this genus in the hot dry valley of the Yuanjiang. *Terminalia franchetii* is an endemic shrub or small tree found in the deep and hot dry river drainages in the Jinshajiang and Yuanjiang. Chloroplast phylogeography and phylogeographic structure of this species revealed that that the modern disjunctive distribution and associated patterns of genetic structure of this species resulted from vicariance caused by several historical drainage capture events, involving the separation of the Jinshajiang from the Yuanjiang [31, 32]. *Tsaiodendron* is a new genus that has been recently found in the hot dry valley of the Yuanjiang [33]. The phylogenetic analyses revealed that it originated in the late Miocene 10.42 Mya and its origin could be correlated with the Red River incision from SW China. These studies on the genera and species reveal that the Jinshajiang separated from the Yuanjiang by river capture events that might have occurred in the Miocene.

More temperate elements are found in the flora of the Jinshajiang, which could be in its present location as part of Himalayas and is at a higher elevation although its deep valley has a typical hot dry climate due to the foehn effect. Its flora was proliferated by temperate elements with the uplift of the Himalayas. The floras of the Yuanjiang (the upper reaches of the Red River) and the Nujiang (the upper reaches of the Salween River) are dominated by tropical elements and the higher ratio of tropical Asian elements could be explained by the events of the southern extrusion of the Indochina Plate and the northward movement of the Burma Plate relative to the Asian Plate to the east [34]. Although the Nujiang is at a similar latitude as the Jinshajiang, its flora is dominated by tropical elements, which could relate to the 1,100 km northward movement of the Burma Plate from its original southern tropic position since the Cenozoic. Our biogeographical studies on the flora of the hot dry valleys in SW China further confirm the geological history of the region.

With physiognomic similarity of the savanna-like vegetation in hot dry valleys of SW China, it has been considered that the savanna-like vegetation of SW China has floristic affinity to the savannas in India and Africa. This is supported by the theory that the Indian Plate drifted away from southern Africa, and moved northward to collide with Eurasia. Here we compare the floristic similarities at the family and generic levels between the flora in hot dry valleys and the floras in the neighboring Gunagxi [35], Myanmar [36], and southern Africa [37], based on available data (Table 7). The comparison shows that the flora of southern Africa has the highest similarity coefficients to the flora of savanna-like vegetations of SW China at the family level (83.95%) and generic level (39.22%) compared to Myanmar and Guangxi Province. This supports the consideration that the savanna-like vegetations of SW China could have floristic affinity to the African savannas during the course of evolution.

**Table 7.** Comparison of floristic similarities at the family and generic levels between the flora in hot dry valleys and floras of the neighboring Gunagxi of SW China, Myanmar, and the southern tropical Africa

| Compared flora | The hot dry<br>valleys (162<br>families, 1038<br>genera) | Guangxi, SW<br>China (209<br>families, 1653<br>genera)<sup>1</sup> | Myanmar (209<br>families, 1903<br>genera)<sup>2</sup> | Southern<br>tropical Africa<br>(224 families,<br>1985 genera)<sup>3</sup> |
| --- | --- | --- | --- | --- |
| Similarity coefficients at family level (%)* |  |  |  |  |
| The dry-hot valleys | 100 |  |  |  |
| Guangxi, SW China | 96.30 | 100 |  |  |
| Myanmar | 96.30 | 94.29 | 100 |  |
| Southern tropical Africa | 83.95 | 77.14 | 77.17 | 100 |
| Similarity coefficients at generic level (%)* |  |  |  |  |
| Dry-hot valleys | 100 |  |  |  |
| Guangxi, SW China | 81.17 | 100 |  |  |
| Myanmar | 79.51 | 73.43 | 100 |  |
| Southern tropical Africa | 39.22 | 31.54 | 33.74 | 100 |

\*Similarity coefficient between A and B = the number of taxa shared by both A and B divided by the
lowest number of taxa of A or B, multiplied by 100%
<sup>1</sup> from Qin & Liu [35]; <sup>2</sup> from Kress et al.[36]; <sup>3</sup> from Leistner [37].

Bucini and Hanan found that woody communities across African savannas were best represented by a sigmoidal response of tree cover to mean annual precipitation, and woody cover increased with rainfall but was modified by disturbances [11]. The savanna-like vegetations in SW China appear to have the same ecology. Bucini and Hanan also found that in addition to climate, other factors, such as soil characteristics, fire, herbivory, and human activities, are important forces in the development and control of vegetation structure and function. Our conclusions are similar in that in addition to climate, fire and human activities are important forces in the development and control of vegetation structure of the savanna-like vegetations of SW China.

## Conclusions

The savanna-like vegetation in southwestern China occurs mainly in deep, hot, and dry valleys of the Jinshajiang (the upper reaches of the Yangtze River), the Yuanjiang (the upper reaches of the Red River), and the Nujiang (the upper reaches of the Salween River) in Yunnan Province. The flora of the vegetation dominates by tropical families and genera except the most species-rich families with cosmopolitan distribution, and is fundamentally tropical in nature.

The geological events since the Cenozoic, could influence the evolution and divergence of the flora in hot dry valleys in SW China. The floras of the Yuanjiang and Nujiang are dominated by tropical elements (77.26% and 74.49% of the total genera, respectively), and have the higher ratio of tropical Asian elements, which could be explained by the events of the southern extrusion of the Indochina Plate and the northward movement of the Burma Plate relative to the Asian Plate to the east, but the flora of the Jinshajiang is composed of half tropical (47.27%) and half temperate (44.96%) elements, which could be influenced by the rapid uplift of the Himalayas and proliferated by temperate elements of the Himalayas, despite being in a deep valley with a typical hot dry climate due to foehn effect. The floristic similarities at the generic and specific levels are higher between the Yuanjiang and Jinshajiang, although these rivers are located a great distance from each other. This supports the theory that the Jinshajiang was once a tributary to the paleo-Red River, i.e., the present Yuanjiang. In the Miocene, disruption of the paleo-drainage occurred via river capture, which led to the separation of the Jinshajiang from the Yuanjiang. Our biogeographical studies on the flora of the hot dry valleys in SW China further confirm the geological history of the region.

Floristic comparison show that the flora of southern Africa has the highest similarity coefficient to the flora in hot dry valleys of SW China. This supports the consideration that the savanna-like vegetations of SW China could have floristic affinity to the African savannas over the course of its evolutionary history.

## Supporting Information

Additional Supporting Information may be found in the online version of this article:

**Appendix S1** Names of the native seed plant families with numbers of the genus and species from the savanna-like vegetations in hot dry valleys, SW China (The circumscriptions of families follow APG III, and ranking alphabetically).

## Acknowledgments

Figures 1 and 2 were made by Yang Jianbo from GIS Lab in Xishuangbanna Tropical Botanical Garden, Chinese Academy of Sciences. We thank Mr. Liu Fangyan from Research Institute of Insect Resources,Chinese Academy of Forestry, offering help in our floristic inventory. TopEdit company (www.topedit.cn)made English polishing for the article. We would like to the thank reviewers’ constructive suggestions on this article.

## Author Contributions

Conceptualization: HZ. Datacuration: LY. Formal analysis: HZ. Funding acquisition: HZ. Investigation: HZ., LY. Methodology: HZ. Project administration: HZ. Resources: HZ., LY. Supervision: HZ. Validation: HZ. Visualization: HZ. Writing-original draft: HZ. Writing-review & editing: HZ.

## Figure legends

Photo 1 Physiognomy of the savanna-like vegetation in the hot dry valleys of Yuanjiang, SW China

Photo 2 Physiognomy of the dry thorny shrubs in the hot dry valleys of Yuanjiang, SW China

